# Normalization by distributional resampling of high throughput single-cell RNA-sequencing data

**DOI:** 10.1101/2020.10.28.359901

**Authors:** Jared Brown, Zijian Ni, Chitrasen Mohanty, Rhonda Bacher, Christina Kendziorski

## Abstract

**Motivation:** Normalization to remove technical or experimental artifacts is critical in the analysis of single-cell RNA-sequencing experiments, even those for which unique molecular identifiers (UMIs) are available. The majority of methods for normalizing single-cell RNA-sequencing data adjust average expression in sequencing depth, but allow the variance and other properties of the gene-specific expression distribution to be non-constant in depth, which often results in reduced power and increased false discoveries in downstream analyses. This problem is exacerbated by the high proportion of zeros present in most datasets.

**Results:** To address this, we present Dino, a normalization method based on a flexible negative-binomial mixture model of gene expression. As demonstrated in both simulated and case study datasets, by normalizing the entire gene expression distribution, Dino is robust to shallow sequencing depth, sample heterogeneity, and varying zero proportions, leading to improved performance in downstream analyses in a number of settings.

**Availability and implementation:** The R package, Dino, is available on GitHub at https://github.com/JBrownBiostat/Dino.

## 1 Introduction

Over the past decade, advances in single-cell RNA-sequencing (scRNA-seq) technologies have significantly increased the sensitivity and specificity with which scientific questions can be addressed (Wu *et al.*, 2017; Hwang *et al.*, 2018; Bacher and Kendziorski, 2016; Haque *et al.*, 2017; Kolodziejczyk *et al.*, 2015). The 10x Genomics Chromium (Zheng *et al.*, 2017) platform, which utilizes a droplet-based, unique-molecular-identifier (UMI) protocol, has become increasingly popular as it provides for rapid and cost effective gene expression profiling of hundreds to tens of thousands of cells.

The use of UMIs has significantly reduced biases due to transcript length and PCR amplification (Tung *et al.*, 2017; Grün *et al.*, 2014; Islam *et al.*, 2014; Zheng *et al.*, 2017). However, technical variability in sequencing depth remains and, consequently, normalization to adjust for sequencing depth is required to ensure accurate downstream analyses (Fig. 1). As a result, a number of normalization methods have been developed to remove the effects of sequencing depth on average expression.

**Figure 1:**
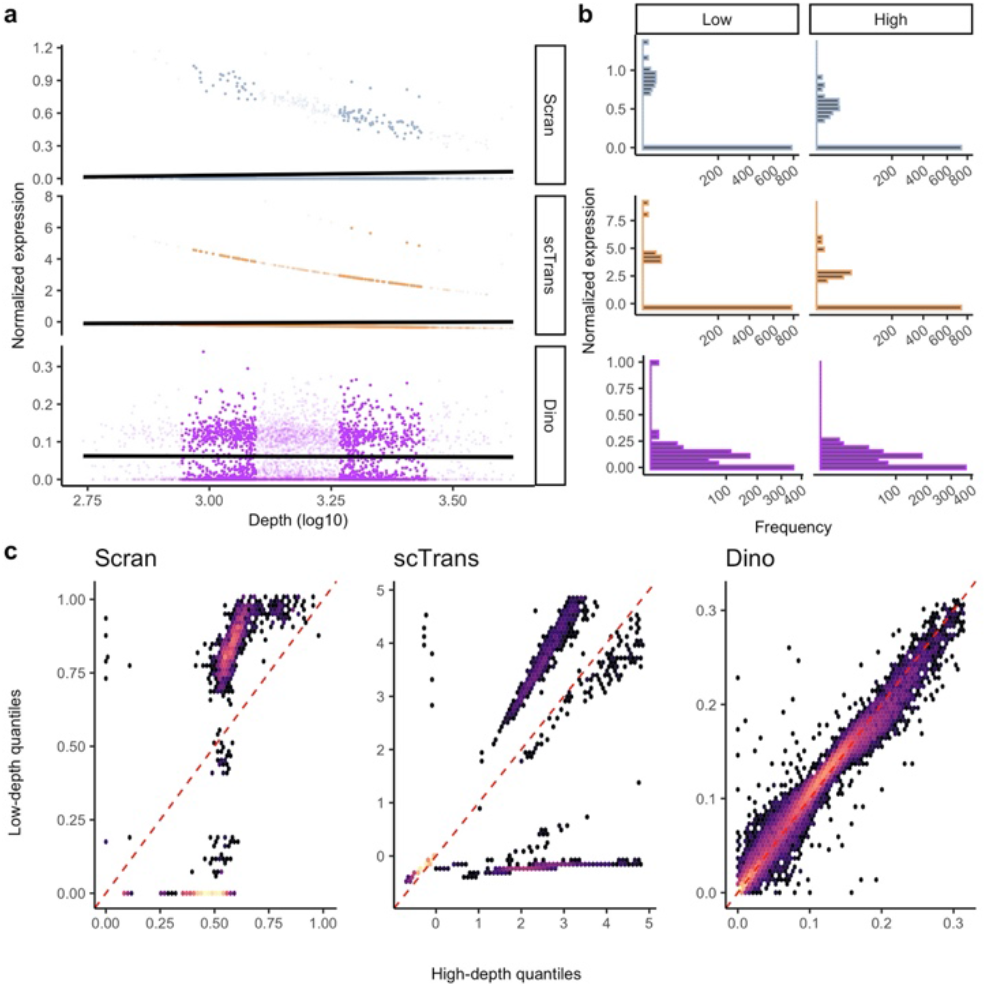
Evaluation of gene-specific expression distributions following normalization. Expression data in the PBMC68K_Pure dataset were normalized by Scran scTransform, and Dino. Normalized expression is shown here for a homogeneous set of cells (CD4+/CD45RO+ memory cells) to minimize the effects of cell subpopulation heterogeneity. a) Normalized expression from a typical gene (NME1) under Scran, scTransform, and Dino plotted against sequencing depth. Fitted regression lines (solid black) show generally constant means across methods. b) Histograms of expression from low and high depth cells (bold in panel a) show that the constant mean is maintained by balancing the changing proportions of zeros, or near zeros in the case of scTransform, with expression shifts in normalized non-zeros. c) Quantile-quantile density plots comparing expression quantiles in the high-depth (x-coordinate) and low-depth cells (y-coordinate) across genes. As in panel b, there are systematic shifts in the distributions.

The earliest methods for normalization were based on global scale factors which adjust all transcripts in a sample (here, a cell) uniformly. In transcripts per ten-thousand (TPT), implemented in the popular Seurat (Butler *et al.*, 2018) pipeline, each transcript within a cell is scaled such that the sum of expression across transcripts within the cell equals ten thousand; transcripts per million (TPM) is similar, but with the target sum equal to one million. Another widely used method, scran (Lun *et al.*, 2016), pools counts across groups of cells to calculate scale factors which are more robust to low sequencing depths.

Bacher *et al*. (Bacher *et al.*, 2017) showed that different groups of genes require different scale factors, which compromises the performance of global scale factor based approaches. To address this, they proposed scNorm which estimates scale factors via quantile regression for groups of genes having similar relationships between expression and depth. While useful, their approach was developed for scRNA-seq data obtained via Fluidigm and similar protocols, and does not apply directly to UMI count data.

Hafemeister and Satija recently demonstrated that analysis of UMI data also requires different distributional parameters for different groups of genes. They approach normalization as a parametric regression problem and introduce scTransform (Hafemeister and Satija, 2019) which models counts using a negative-binomial generalized linear model (glm). In scTransform, parameter estimates are smoothed across genes such that genes with similar average expression also have similar model parameters. Normalized data is then given by Pearson residuals from the regression and, as a result, the normalized expression of a typical gene has mean zero and unit variance. The approach attenuates the dependence of both the mean and variance on sequencing depth, but maintains many of the depth-dependent properties of the normalized expression distribution (Fig. 1), which impacts downstream analyses (Fig.2).

**Figure 2:**
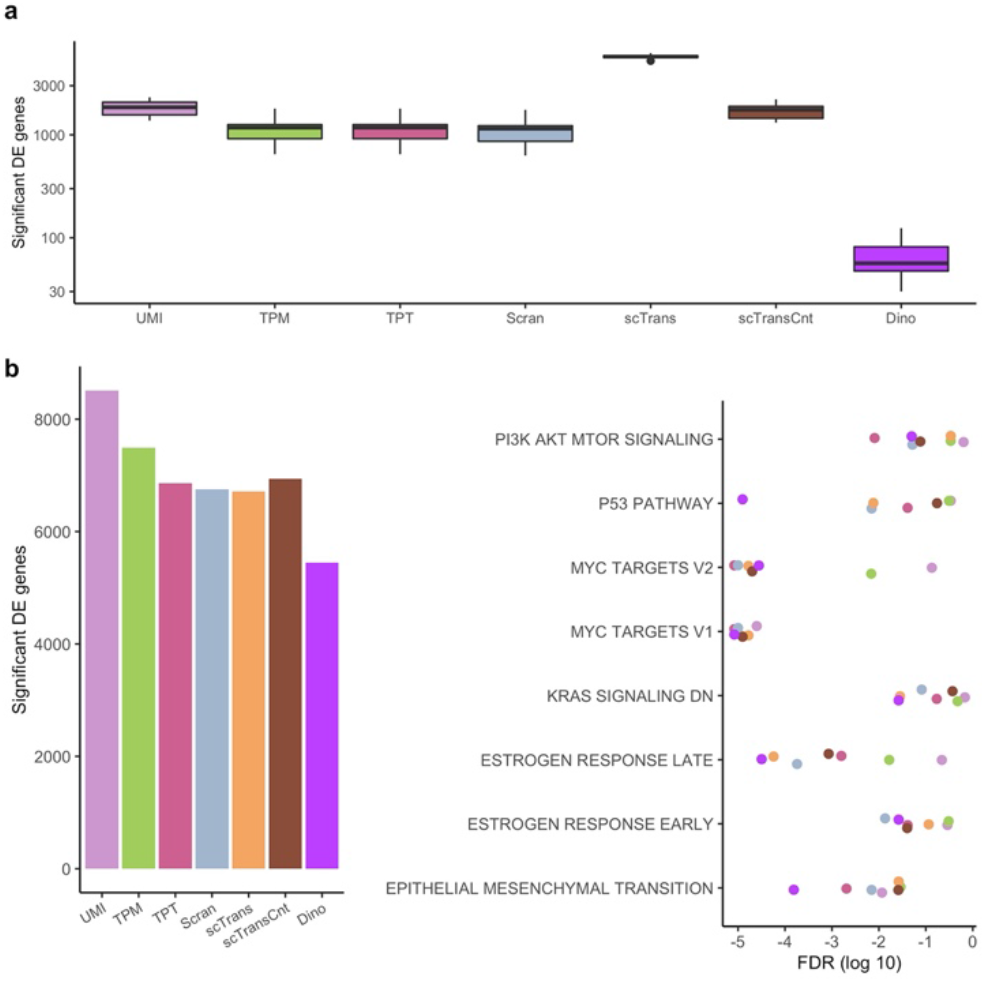
The effects of normalization on downstream DE and enrichment analysis. a) Expression data from the PBMC68K_Pure dataset were normalized and genes were tested for DE using a Wilcoxon rank sum test between low-depth and high-depth cells within cell-type annotations. Box plots show DE genes; given cells only differ in depth, DE identifications are considered false positives. b) Expression data from the EMT dataset were analyzed using Monocle2 to identify genes with significantly variable expression over pseudo-time. (Left) Total numbers of significant genes. (Right) Significance values of enriched GO terms, colored for each normalization method, for the subset of terms previously identified as defining expression shifts during endothelial mesenchymal transition.

To address this, we present Dino, an approach that utilizes a flexible mixture of negative binomials model of gene expression to reconstruct full gene-specific expression distributions which are independent of sequencing depth. By treating zeros as expected values, the negative binomial components are applicable to shallow sequencing. Additionally, the mixture component is robust to cell heterogeneity as it accommodates multiple centers of gene expression in the distribution. By directly modeling (possibly heterogenous) gene-specific expression distributions, Dino outper-forms competing approaches, especially for datasets in which the proportion of zeros is high as is typical for modern, UMI based protocols.

## 2 Methods

Our proposed method for <u>di</u>stributional <u>nor</u>malization, Dino, reconstructs gene-specific expression distributions and provides normalized estimates of expression by constrained sampling from those distributions. Specifically, Dino assumes a Poisson model on observed counts and models the distribution of Poisson means as a mixture of Gammas across cells, conditioned by sequencing depth. This Gamma-Poisson model is equivalent to modeling counts as a mixture of Negative Binomials. Normalized expression is then sampled from cell-specific posterior distributions. The estimated distributions are constructed across cells and approximate the flexibility of a non-parametric approach in order to accommodate varying degrees of heterogeneity in the cell populations under study.

### 2.1 Statistical model

The count data produced by UMI sequencing protocols lend themselves naturally to a glm parameterized by sequencing depth (Anders and Huber, 2010; Hafemeister and Satija, 2019); and the random sampling of barcoded molecules from a large pool for sequencing is theoretically well modeled by independent Poisson distributions on each gene (Townes *et al.*, 2019). Furthermore, Poisson means are expected to scale proportionally with sequencing depth (Anders and Huber, 2010; Lun *et al.*, 2016; Hafemeister and Satija, 2019; Townes *et al.*, 2019), giving counts *Y_gj_* from gene *g* in cell *j* the distribution *Y_gj_~Pois(λ_gj_δ_j_)*. Defining *δ_j_* to be the cell-specific sequencing depth, *λ_gj_* then represents the latent level of expression for gene *g* in cell *j*, corrected for depth. Note that *λ_gj_* is cell dependent given that latent levels of expression for a gene may vary across cells due to population heterogeneity. For convenience of interpretation on the *λ_gj_*, estimated depths are scaled such that *δ_jMed_=1* where *jMed* indexes the median depth cell by default.

The Dino algorithm defines the distribution of *λ_gj_* across cells to be the gene-specific expression distribution of interest. If one assumes a Gamma distribution on the *λ_gj_*, then the marginal distribution of the *Y_gj_* is Negative Binomial, the model assumption made by scTransform and DESeq2, as well as the dimension reduction method, ZINB-WaVE (Risso *et al.*, 2018), and others. However, to increase the precision with which the full gene-specific expression distribution may be estimated, Dino further assumes the *λ_gj_* arise from a mixture of Gamma distributions, and defines normalized expression as samples from the posterior distribution of the *λ_gj_*.

As distribution estimation and normalization are both performed at the gene level, the gene subscript is hereafter dropped, and it is noted that computations are repeated across genes. This defines the final model as:

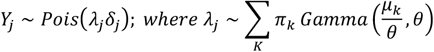

for shape *μ_k_/θ*, scale *θ*, and such that *Σ*_*k*_*π*_*k*_=*1*. This parameterization is chosen to define the distribution in terms of its mean, *μ_k_. K* is chosen to be sufficiently large to accommodate both cellular heterogeneity and within-cell-type over-dispersion of the *λ_j_* with respect to a single *Gamma(s_k_,θ)* mixture component. By default, *K* is the minimum of 200 and the square root of the number of *Y_j_* which are greater than zero. Since the negative binomial distribution can be alternately defined as the Gamma-Poisson distribution, this formulation has the reassuring additional interpretation of defining *Y_j_* as a mixture of Negative Binomials.

### 2.2 Sampling normalized values from the posterior

The posterior distribution on *λ_j_* is straightforward to compute:

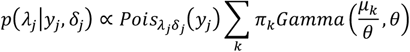

which reduces to

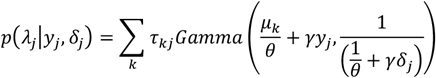

where *τ_kj_* is the conditional likelihood that *λ_j_* belongs to component *k* given *y_j_* and *δ_j_*, and *γ* is a concentration parameter. Our testing (not shown) has demonstrated that strict resampling from the posterior distribution can lead to excessive variance which can obscure biological features of interest. Therefore, *γ* is added to both reduce the normalized variance and center normalized values around their corresponding scale-factor variants:

Default values of *γ*=*25* have proven successful. This adjustment can be seen as a slight bias in the normalized values towards a scale-factor version of normalization, since, in the limit of *γ*, the results converge to y_*j*_/δ_*j*_. A modified expectation-maximation (EM) algorithm (Jamshidian and Jennrich, 1997), described in detail in Supplementary Section S1.1, is used to estimate *μ_k_*. Separate values of *θ* are estimated for each gene based on gamma kernel density estimation (Chen, 2000) as described in Supplementary Section S1.2. Adjustments to *δ_j_* to accommodate slight deviation from strict expression scaling with depth are described in Supplementary Section S1.3. Finally, parameter initialization relies on a modified application of quantile regression (Powell, 1984, 1986; Branham, R. L., 1982) and is described in Supplementary Section S1.4.

### 2.3 Datasets

Results from six publicly available datasets are evaluated: PBMC68K_Pure, PBMC5K_Prot, MaltTumor10K, MouseBrain, PBMC68K, and EMT. Where applicable, analyzed expression was derived from unfiltered gene-barcode matrices, with empty droplets removed by the tools in the R package DropletUtils (Lun *et al.*, 2018).

PBMC68K_Pure is a partner dataset to PBMC68K (Zheng *et al.*, 2017) produced by fluorescence activated cell sorting (FACS) of peripheral blood mononuclear cells (PBMCs) into 10 cell types and separately sequencing each group. One group was then computationally separated into two resulting in 11 annotated cell-types. These cell-type annotations are considered here as ground truth when evaluating the effects of normalization on downstream clustering, and for increased accuracy, the six most homogenous cell-types from visual inspection of tSNE plots (van der Maaten and Hinton, 2008; Van Der Maaten, 2014) were subset: CD4+ T Helper2, CD4+/CD25 T Reg, CD4+/CD45RA+/CD25- Naive T, CD4+/CD45RO+ Memory, CD56+ NK, and CD8+/CD45RA+ Naive Cytotoxic.

EMT is a dataset of 5,004 MCF10A mammary epithelial cells induced to undergo spontaneous endothelial to mesenchymal transitions (EMTs) through the cellular detection of neighboring unoccupied space (McFaline-Figueroa *et al.*, 2019). This spatial effect allowed the authors to dissect an inner region *a priori* expected to be primarily endothelial cells and an outer region *a priori* expected to be primarily mesenchymal cells which were then sequenced separately. From all the data published by the authors, the EMT dataset we consider is denoted “Mock” in the barcode metadata. Included in the initial publication, the authors describe eight gene sets from the Hallmark collection (Liberzon *et al.*, 2015) which they consider to be significantly enriched for activity during EMT. We take this set of terms as a ground truth for assessing power of each normalization method.

Results from the PBMC68K_Pure and EMT datasets are shown in the manuscript while results from other datasets are provided in the supplement unless otherwise noted. Details on each of the datasets as well as their pre-processing are provided in Supplementary Section S2.

The case study datasets are also used to generate simulated datasets with expression profiles designed to closely mirror experimentally observed cells. In brief, for a given case study data set, unsupervised clustering is used to define clusters. Two cells with similar sequencing depths are sampled from a cluster and expression is summed across the two cells to make a pseudo-cell. Expression from this pseudo-cell is then down-sampled using a binomial distribution to generate two new simulated cells which differ in sequencing depth, but otherwise have equivalent expression (EE) across all but 10 genes for which differential expression (DE) is induced. This process is repeated to generate a collection of cells with the same set of EE and DE genes; and the process is repeated again for other clusters to simulate subpopulation heterogeneity. These simulated datasets are then used to quantify power and false positive rates for DE testing following different normalization methods. Full simulation details are described in Supplementary Section S3.

### 2.4 Application of normalization methods

For each dataset considered, normalized estimates of expression were obtained from Dino (v0.6.0), scran (v1.16.0), scTransform (v0.2.1), TPM, and TPT. We also consider un-normalized UMIs for reference. Further information on package defaults, annotation versions, and other software is given in Supplementary Section S4. The implementation of scTransform provides both normalized expression in terms of regression residuals (recommended for most analysis applications by the authors) and normalized expression in terms of a corrected UMI counts matrix. We consider both in this manuscript and refer to the residuals matrix as scTrans and the counts matrix as scTransCnt.

### 2.5 Evaluation of gene-specific expression distributions

To evaluate the extent to which a given normalization method removes the effect of sequencing depth on expression, we use linear regression to quantify the relationship between expression and depth. We also compare the gene-specific expression distributions between low-depth cells (depth between the 5% and 25% quantiles) and high-depth cells (depth between the 75% and 95% quantiles).

While such a comparison can be done visually for individual genes, to evaluate the extent to which there are shifts in these distributions across genes, we construct QQ plots aggregated across multiple genes. Within a cell type annotation, we sample genes from the bottom 90% of expression (geometric mean of un-normalized UMIs), omitting only the high expressors which perform similarly across normalization methods. For each gene, we compute a grid of quantiles from the cells in the low-depth group (defined above) and a corresponding grid of quantiles from the cells in the high-depth group. These vectors of quantiles define points on the QQ plot with the high-depth quantiles forming the x-coordinate and the low-depth quantiles defining the y-coordinate. Application of this procedure across genes builds a larger, representative set of points on the QQ plot which are then transformed into densities for visualization.

### 2.6 Identification of differentially expressed genes and evaluation of clusters

Genes that are DE between two groups are identified using the Wilcoxon rank sum test, which is the default in the Seurat pipeline at the time of writing. We also consider MAST (v1.14.1) (Finak *et al.*, 2015) as implemented in Seurat (v3.2.0). For each method, we define a list of genes with false discovery rate less than 1%. Specifically, a gene is defined as DE if its Benjamini and Hochberg adjusted p-value is less than 0.01 (Benjamini and Hochberg, 1995).

To identify genes that are DE over pseudo-time in the EMT data, we use a simplified analysis approach similar to that of the original authors. Highly variable genes define a pseudo-time ordering of cells using Monocle2 (v2.16.0) (Trapnell *et al.*, 2014; Qiu *et al.*, 2017). DE genes are then defined as genes with a significant trend over pseudo-time. Significance is determined via a likelihood ratio test between a natural spline regressed against pseudo-time and an intercept only null model, implemented in Monocle2. As with the two-group case, we define a gene as DE if its Benjamini and Hochberg adjusted p-value is less than 0.01. Full details are provided in Supplementary Section S4.

To perform clustering analysis, clusters are estimated using 25,000 cells selected at random from each dataset; the same set of 25,000 cells is used for normalization by each method. After normalization, the top 1,000 highest variance genes are reduced to 25 principal components on the log scale with a +1 psuedo-count for all methods to stabilize the variance except for scTrans as it performs a similar variance stabilization internally. Louvain graph-based clustering, implemented in Seurat, is used to identify clusters. Cluster accuracy is quantified by computing the adjusted Rand index (ARI) between discovered clusters and annotations, and clusters are visualized using tSNE dimension reduction.

Clustering analysis is also performed in an environment of exaggerated differences in sequencing depth where half of the original 25,000 cells sampled from each dataset are randomly selected and down sampled to 25% of their original depth. Normalization, clustering, and calculation of ARI are then performed as previously descried across all 25,000 cells. Full details are provided in Supplementary Section S4.

## 3 Results

### 3.1 Depth-dependent patterns in normalized data

To compare Dino, scran, scTransform, TPM, and TPT, we normalized the PBMC68K_Pure data using each method. Given the purification done with FACS, cells within a given annotation should be largely homogeneous and, consequently, should exhibit little difference in expression among cells. Examination of the normalized expression between low-depth and high-depth CD4+/CD45RO+ memory cells shows that existing methods exhibit significant depth-dependent effects. As shown in Fig.1a, for a normalized gene to maintain average expression that is relatively independent of depth, the higher proportion of zeros in the low-depth group is balanced by inflation of the non-zeros. This leads to shifts in the densities of the normalized non-zeros between low and high-depth cells, as shown in Fig.1b. To evaluate this effect across all genes, we compare the quantiles in low-depth and high-depth cells. Figure 1c. demonstrates systematic shifts in the normalized expression distributions. Normalized expression from Dino mitigates these effects, producing more equivalent expression distributions across low and high depth cells. Similar results are shown for other case study datasets in Supplementary Figure S1.

### 3.2 Effects of normalization on differential expression analysis

To evaluate the extent to which depth-dependent differences in the expression distributions affect downstream differential expression (DE) analysis, we used the Wilcoxon rank-sum test to identify DE genes between low and high-depth cells within each of the 6 annotated cell-types in the PBMC68K_Pure dataset, resulting in 6 separate measures of DE genes. Fig. 2a demonstrates that many DE genes are identified by most methods. As each cell-type is expected to be homogeneous, these identifications are considered false positives. Similar results are shown in Supplementary Figure S2 for the MaltTumor10K and PBMC5K_Prot datasets. These datasets were considered as they have pseudo-annotations available (see Supplementary Section S2 for dataset details).

To evaluate DE analysis in the positive case where DE genes are expected to exist, we consider a case study dataset of cells undergoing spontaneous endothelial to mesenchymal transitions (EMT dataset) (McFaline-Figueroa *et al.*, 2019). Cells along the primary branch of the differentiation tree are tested for DE, here defined as having a significant change in expression over pseudo-time.

As with the negative control examples of Fig. 2a and S2, Dino discovers the fewest DE genes (Fig. 2b, left). Results from an enrichment analysis suggest that the reduction in the number of DE genes found by Dino is likely due to a reduction in false positives, as with the negative control, rather than an undesirable reduction in power. Specifically, we performed gene set enrichment analysis (GSEA) on the hallmark collection of gene sets (Liberzon et al., 2015). Enrichment significance values are plotted in the right panel of Fig. 2b for the set of terms identified by McFaline-Figueroa et al. as significant markers of EMT activity. Dino normalization results in competitive significance in 3 of the 8 terms and shows the highest GSEA significance in 5 of the 8. Notably, Dino shows improved enrichment results for the term defining endothelial mesenchymal transition; the Dino adjusted p-value (1.5e-4) is more than an order of magnitude more significant than the nearest alternate method (TPT, adjusted p=2.0e-3).

Simulated datasets provide further insights consistent with the case study results regarding DE. Specifically, we simulated heterogenous cell populations from experimentally observed expression with known DE and EE genes (see Supplementary section S3 for simulation details). Following normalization, DE genes are identified between cluster pairs by a Wilcoxon rank sum test. Fig. 3 plots average ROC curves for each normalization method where the average is taken over 50 datasets simulated from the PBMC68K_Pure dataset. Average true positive rates (TPR / Power) and false positive rates (FPR) are given in Supplementary Table S1 for simulations based on other case study datasets.

**Figure 3:**
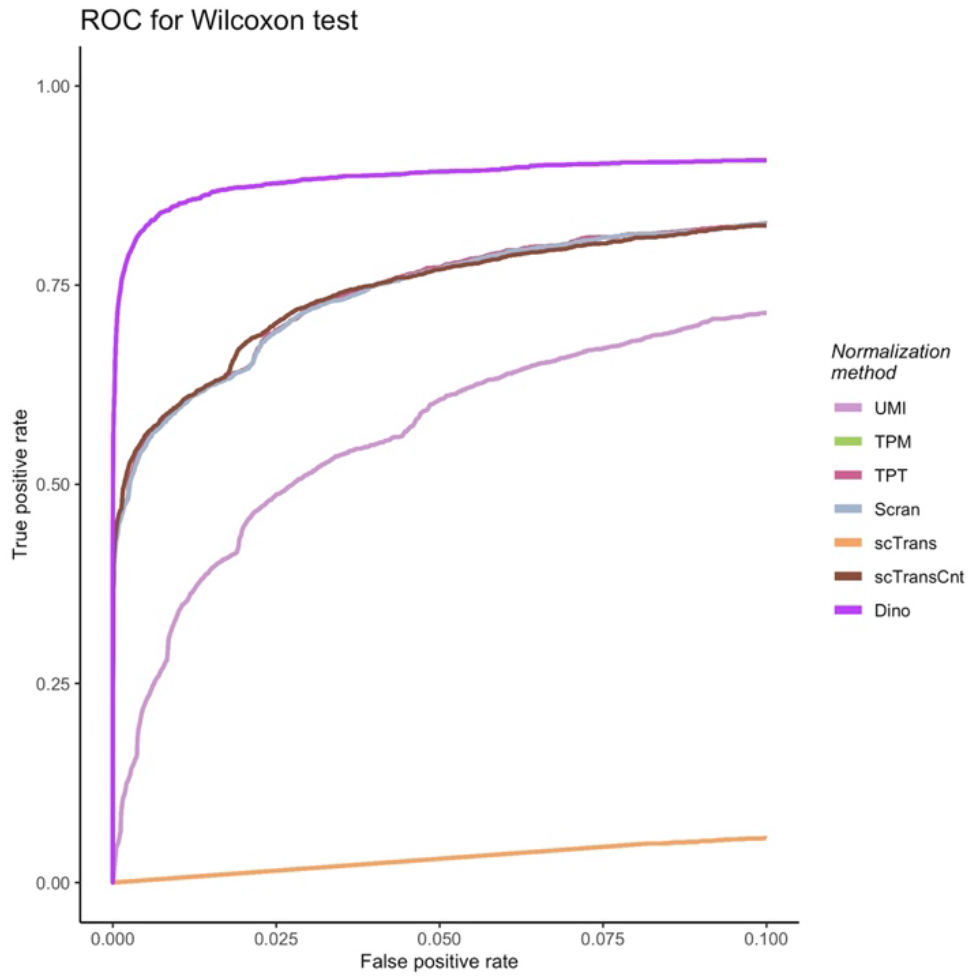
The effects of normalization on downstream DE analysis. Simulated data based on the PBMC68K_Pure dataset were normalized using each method. ROC curves colored by normalization method define the relationship between average TPR and average FPR for a Wilcoxon rank sum test, where the average is calculated across 50 simulated datasets.

In most cases it can be observed that high power is confounded by high FPRs. As with the negative control, however, Dino controls FPR to much lower levels. ROC curves across datasets are considered in Supplementary Figures S3 and S4, and Power and FPR for MAST tests are described in Supplementary Table S2.

### 3.3 Effects of normalization on clustering

The effect of normalization on clustering was evaluated by comparing clusters derived using data normalized by each method with the annotations in the PBMC68K_Pure dataset using the adjusted Rand index (ARI). As shown in Fig. 4 for one sub-sample of cells, Dino, scran, and scTrans outperform other methods; Dino shows slightly although not significantly improved performance over scTrans and scran (Fig. 4a). To exacerbate the differences in sequencing depth, we randomly down-sampled half of the 25000 cells to 25% of their original sequencing depth. Fig. 4b shows the derived clusters with the down-sampled cells plotted bold compared to the unmodified cells. Some effect of depth is observed in the clustering, and ARIs for all normalization methods decreased, as expected. However, Dino normalized data retains a more accurate differentiation between cells with an ARI of 0.480 compared to 0.357 and 0.350 for scTrans and scran, respectively.

**Figure 4:**
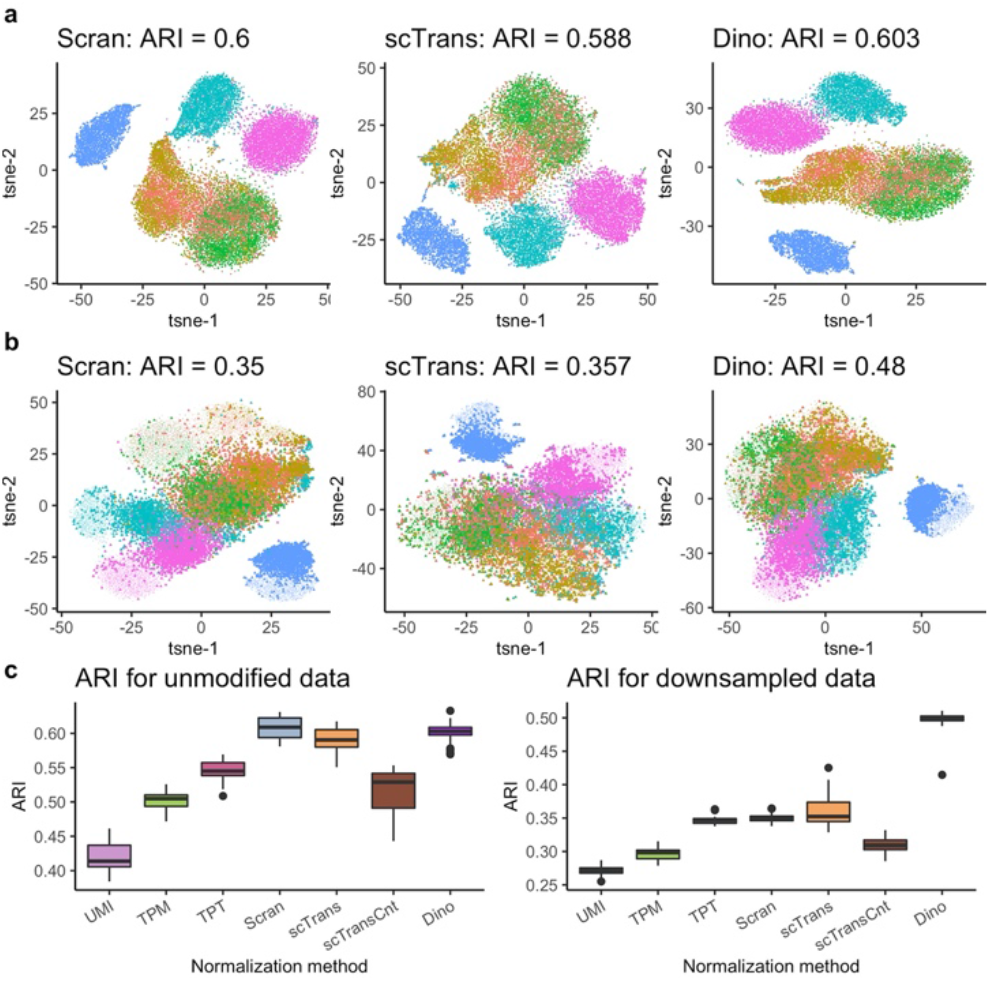
The effects of normalization on clustering. a) tSNE plots of normalized PBMC68K_Pure data, colored by 6 cell-type annotations, show similarly high accuracy between methods. b) The same clustering plots as in (a), but with half the data down-sampled prior to normalization to produce greater differences in sequencing depths. c) Boxplots of ARIs for multiple un-modified and down-sampled datasets.

Fig. 4c shows results from the analysis repeated across multiple samples of 25,000 cells. As with the sample shown in Figs. 4a, Dino and scran perform comparably (medians of 0.601 and 0.609 respectively), and uniformly better than scTrans (median of 0.591). In the down-sampled case, Dino performs significantly better than competing methods (p<2.2e-16 under a t-test). Similar results are shown for the MaltTumor10K and PBMC5k_Prot datasets (Supplementary Figures S5 and S6).

## 4 Discussion

The 10x platform and similar protocols provide unprecedented cellular resolution in expression profiling. However, the use of UMIs does not remove the need for effective normalization in the analysis of such data, and the extreme sparsity of these high-throughput experiments introduces new challenges for depth correction. Dino adapts to these challenges by correcting the entire expression distribution of each gene in depth, rather than only correcting mean expression as with most existing methods. This increases both power and precision in down-stream analysis.

Dino normalizes observations by resampling from gene-specific expression distributions, conditional on observed expression and depth. This resampling accommodates the varying proportions of zeros across depth induced, in large part, by technical artifacts. In addition, by resampling from the full gene-specific distribution, Dino produces greater homogeneity of normalized expression across sequencing depth within cell type, and therefore leads to more accurate downstream analyses that are robust to heterogeneous cell populations.

## Acknowledgements

The authors would like to thank Matt Bernstein for conversations that improved the manuscript.

## References

Anders,S. and Huber,W. (2010) Differential expression analysis for sequence count data. Genome Biol., 11, R106.

Bacher,R. et al. (2017) SCnorm: Robust normalization of single-cell RNA-seq data. Nat. Methods, 14, 584–586.

Bacher,R. and Kendziorski,C. (2016) Design and computational analysis of singlecell RNA-sequencing experiments. Genome Biol., 17, 1–14.

Benjamini,Y. and Hochberg,Y. (1995) Controlling the False Discovery Rate: A Practical and Powerful Approach to Multiple Testing. J. R. Stat. Soc. Ser. B, 57, 289–300.

Branham, R. L. J. (1982) Alternatives to least squares. Astron. J., 87, 928.

Butler,A. et al. (2018) Integrating single-cell transcriptomic data across different conditions, technologies, and species. Nat. Biotechnol., 36, 411–420.

Chen,S.X. (2000) Probability Density Function Estimation Using Gamma Kernels. Ann. Inst. Stat. Math., 52, 471–480.

Finak,G. et al. (2015) MAST: A flexible statistical framework for assessing transcriptional changes and characterizing heterogeneity in single-cell RNA sequencing data. Genome Biol., 16, 1–13.

Grün,D. et al. (2014) Validation of noise models for single-cell transcriptomics. Nat. Methods, 11, 637–640.

Hafemeister,C. and Satija,R. (2019) Normalization and variance stabilization of single-cell RNA-seq data using regularized negative binomial regression. Genome Biol., 20, 296.

Haque,A. et al. (2017) A practical guide to single-cell RNA-sequencing for biomedical research and clinical applications. Genome Med., 9, 1–12.

Hwang,B. et al. (2018) Single-cell RNA sequencing technologies and bioinformatics pipelines. Exp. Mol. Med.,50.

Islam,S. et al. (2014) Quantitative single-cell RNA-seq with unique molecular identifiers. Nat. Methods, 11, 163–166.

Jamshidian,M. and Jennrich,R.I. (1997) Acceleration of the EM Algorithm by using Quasi-Newton Methods. J. R. Stat. Soc. Ser. B (Statistical Methodol., 59, 569–587.

Kolodziejczyk,A.A. et al. (2015) The Technology and Biology of Single-Cell RNA Sequencing. Mol. Cell, 58, 610–620.

Liberzon,A. et al. (2015) The Molecular Signatures Database Hallmark Gene Set Collection. Cell Syst., 1, 417–425.

Lun,A. et al. (2018) Distinguishing cells from empty droplets in droplet-based single-cell RNA sequencing data. bioRxiv, 234872.

Lun,A.T.L. et al. (2016) Pooling across cells to normalize single-cell RNA sequencing data with many zero counts. Genome Biol., 17, 1–14.

Van Der Maaten,L. (2014) Accelerating t-SNE using tree-based algorithms. J. Mach. Learn. Res., 15, 3221–3245.

van der Maaten,L. and Hinton,G. (2008) Visualizing High-Dimensional Data Using t-SNE. J. Mach. Learn. Res., 9, 2579–2605.

McFaline-Figueroa,J.L. et al. (2019) A pooled single-cell genetic screen identifies regulatory checkpoints in the continuum of the epithelial-to-mesenchymal transition. Nat. Genet., 51, 1389–1398.

Powell,J.L. (1986) Censored regression quantiles. J. Econom., 32, 143–155.

Powell,J.L. (1984) Least absolute deviations estimation for the censored regression model. J. Econom., 25, 303–325.

Qiu,X. et al. (2017) Reversed graph embedding resolves complex single-cell trajectories. Nat. Methods, 14, 979–982.

Risso,D. et al. (2018) A general and flexible method for signal extraction from single-cell RNA-seq data. Nat. Commun., 9, 284.

Townes,F.w. et al. (2019) Feature selection and dimension reduction for single-cell RNA-Seq based on a multinomial model. Genome Biol., 20, 1–16.

Trapnell,C. et al. (2014) The dynamics and regulators of cell fate decisions are revealed by pseudotemporal ordering of single cells. Nat. Biotechnol., 32, 381–386.

Tung,P.Y. et al. (2017) Batch effects and the effective design of single-cell gene expression studies. Sci. Rep., 7, 1–15.

Wu,A.R. et al. (2017) Single-Cell Transcriptional Analysis. Annu. Rev. Anal. Chem., 10, 439–462.

Zheng,G.X.Y. et al. (2017) Massively parallel digital transcriptional profiling of single cells. Nat. Commun., 8, 1–12.

