## Supplementary materials for "Normalization by distributional resampling of high throughput single-cell RNA-sequencing data"

### S1 Dino algorithm – additional details

#### S1.1 EM iterations

To speed computation, the distribution on the *λ_j_* is approximated by a mixture of point masses: *λj~Σ_k_π_k_μ_k_*. The expectation maximization (EM) algorithm then has fast, closed form iterations for estimating *π_k_* and *μ_k_*, which are further accelerated using a quasi-Newton adjustment to the EM update (Jamshidian and Jennrich, 1997). In brief, this quasi-Newton adjustment replaces the usual EM update, *g(τ)*, where *τ=[π, μ]*, with a “corrected” EM step: *g(τ)-Sgl(τ)* where *S* is an estimate of the inverse Hessian of the likelihood function based on the Broyden-Fletcher- Goldfarb-Shanno symmetric rank 2 update and *gl(τ)* is the gradient of the likelihood function. In our application, the default step length of 1, applied to the entire corrected EM step, is adjusted as necessary to accord with the strong Wolfe conditions.

The EM algorithm requires initialization of the means *λ_k_* and the cluster occupancy rates *π_k_*. For simplicity, the rates *π_k_* are initialized as *1/K*, and the *λ_k_* are initialized both to remove unnecessary iterations in the model fitting and to conform with these initial rates. To accomplish this, we derive the starting *λ_k_* values from equal spacings on an estimate of the CDF of the *y_j_* at unit depth (*δ=1*). Details of estimating the CDF are provided below in Section S1.4.

#### S1.2 Dispersion parameter estimation

After model fitting, a Gamma kernel density estimator is used to compute the estimated mixture of gamma components based on the fitted *μ_k_* with weights *π_k_*. In particular, given an observed sample *{μ_k_}*, the underlying mixture of Gammas can be estimated as *λ_j_~Σ_k_π_k_Gamma(μ_k_/θ,θ)* (Chen, 2000) where *θ* is estimated by bootstrapping kernel bandwidth estimates for which the resampling is weighted by the *π_k_*.

#### S1.3 Depth adjustment

Sequencing depth for cell *j*, *δ_j_*, is typically estimated as the sum of UMIs across genes for a given cell, or more generally, as the sum of expression alignment estimates for a given sample. Given this, a cell with zero depth should have zero counts, and a glm fitted to expression vs. depth should have a zero intercept and linear relationship with depth, which corresponds to a slope coefficient of 1 under the usual log link function. Such values, however, are not universally observed, although calculated coefficients are generally close to 1 (Bacher *et al.,* 2017; Hafemeister and Satija, 2019). Note that, as mentioned in the main text, in our application depths are scaled such that the median depth *δ_jMed_=1*.

To obtain improved estimates of *δ_j_*, we estimate the slope (under the log link-function) between counts and depth. Specifically, we randomly sample a subset of genes (10,000 by default), with sampling weighted by the inverse density of the gene expression level to ensure proper representation of the (relatively rare) high expressing genes. To accommodate sub-population heterogeneity, we perform clustering on these data using the methods implemented in scran (Lun et al., 2016), and fit a Poisson glm to each gene, generating a unique slope estimate for each gene with each cluster of cells getting a unique intercept. A dataset-wide slope *s* is computed as the mean of fitted slopes to each gene; *s* is close to 1 for most datasets. Depths *δ_j_* are then corrected prior to use in the above mixture model as *δ_j_’=sδ_j_-(s-1)*. To avoid the rare occurrence of negative depths post-transformation, *s* is bounded above and below such that the minimum transformed depth is at least *exp(-3)* its original value.

#### S1.4 Estimation of the gene-specific CDF by cLAD regression

This eCDF in turn can be efficiently calculated for each gene by a modified application of censored least absolute deviations (cLAD) regression (Powell, 1984, 1986) at a carefully chosen grid of given intercepts. cLAD regression has the benefit here of solving for linear functions of quantiles in data – hence its alternate name, quantile regression. The pairing of fitted intercepts (quantiles) and corresponding percentiles then define points along the desired eCDF.

Regression calculations are conducted on the log-log scale, so the both the depths *δ_i_* and the counts *y_j_* are log transformed with a floor of *log(0.999)* in the case of the observed counts; *d_j_=log(δ_j_)*, *z_j_=max{log(y_j_), log(0.999)}*. The implicit addition of a *≈1* pseudo count to the observed zeros prior to log transformation is motivated by the fact that computations are performed using censored quantile regression, which can be written as standard quantile (LAD) regression on the subset of data to the right of the intersection of the regression curve and censoring threshold (Powell, 1986). In particular, the regression model for the observed data is:

$$z_{j}=\max\left\{ \log\left( 0.999 \right),\beta_{0}+d_{j}+\epsilon_{j} \right\}$$

where *β_0_* is some gene-specific intercept and *ϵ_j_* is a random error term following some, possibly complex distribution. For example, in the presence of population heterogeneity, the *ϵ_j_* might follow a bi-modal distribution. Placing the zeros at *log(0.999)* allows the regression solutions to be defined in terms of the observed data as the censoring threshold is placed just below log(1). The choice of a constant slope term (the implicit coefficient of 1 on the depth term), while mathematically convenient in the following, is also not unreasonable. On the multiplicative (log) scale, expression should have about a slope one relationship with depth across all genes, recalling the above comment that both counts and depth are log-transformed for the cLAD regression.

**Estimation of expression distribution quantiles:** The empirical cumulative distribution functions (eCDFs) are denoted by vectors of quantiles and the associated estimated percentiles. The eCDFs are independent of depth insofar as they are location families in depth. The specific eCDF to estimate for each gene will be the one at the “target depth” to which the data are to be normalized. This target depth is the median depth of the observed data, which is centered at 0 (log scale, or 1 on the count scale). To calculate the sample quantiles and percentiles of the eCDF, and as a natural extension of the censored linear model of the data distribution, the mathematics of censored least absolute deviations regression (cLAD) are adopted. The cLAD minimization problem is

$$\min_{\beta_{0}} \left\{ \sum_{j} p_{\pi}\left[ z_{j}-\left( \beta_{0}+z_{j} \right)^{*} \right] \right\}$$

where *p_π_(x)=[π-𝕀(x<0)]*x*, *(x)^*^=max{log(0.999), x)*, and *π* is some percentile (eg. *π=0.5* for median quantile regression). It has previously been shown that cLAD regression has beneficial properties, particularly consistency in the presence of only weakly defined residual distributions (Powell, 1984, 1986). Additionally, study of the solution set to cLAD regression has shown that solution coefficients define lines which pass through at least as many data points as there are free parameters (Branham, R. L., 1982). Given this, the regression problem can be significantly reduced. Given a fixed percentile, *π*, the set of possible solutions is confined to a set of parallel lines – one intersecting each point in the data set – which is finite for finite data. The regression problem is then to determine which line, uniquely defined by its intercept *β_0_*, minimizes the loss function.

As such, the set of quantiles defining the eCDF is easily defined. In particular, the eCDF quantiles are the set of intercepts, *β_0j_*, defining the lines passing through each of the data points. The intercepts/quantiles refer to the previously mentioned “grid of given intercepts.” Since this set of intercepts contains the minimizing solution regardless of the choice of *π* in the regression problem, any additional quantiles would by definition be redundant. Specifically, the set of quantiles *{q_j_}* is defined as *{q_j_}={β_0j_}={z_j_-d_j_}*.

**Estimation of expression distribution percentiles:** For simplicity, consider a highly expressed gene for which there are no observed zero counts prior to log transformation and suppose without loss of generality that the quantiles *{q_i_}* are all unique and that the indices *j* are of decreasing order such that *q_j_>q_j+1_∀j*, *q_1_=max{q_j_}*. In this case, computing the percentiles associated with each quantile is trivial:

$$p_{j}=\frac{1}{n}\sum_{i} \mathbb{I}\left( q_{i}\leq q_{j} \right)=\frac{n-j+1}{n}$$

where *n=|{q_j_}|*. The more general case where a gene may contain zeros, possibly many zeros, is more complicated. However, given the assumption that the residual distribution (*ϵ_j_*) is constant in depth, a simple and analogous solution exists:

$$p_{j}=\frac{n_{j}-\left| \left\{ q_{i} \right\}_{j} \right|}{n_{j}}$$

where

$$\Lambda_{j}=\left\{ i:\delta_{i}\geq\delta_{j}-\left( z_{j}-\log\left( 0.999 \right) \right) \right\}$$

$$n_{j}=\left| \Lambda_{j} \right|$$

$$\left\{ q_{i} \right\}_{j}=\left\{ q_{i}:q_{i}>q_{j}, i\in\Lambda_{j} \right\}$$

The linear interpolation of the set $\left\{ \left( q_{j},p_{j} \right) \right\}$defines an eCDF from which the *λ_j_* can be initialized.

**Monotonicity correction:** In practice, this estimate for the percentiles *p_j_* can become unstable for high *j* (when *n_j_* becomes small). Additionally, some forms of population heterogeneity can cause estimation problems, especially as they can violate the assumption of constant residual distribution. Here we derive the above formulation for the percentiles as well as corrections for these situations.

Recall the example of the highly expressing gene mentioned above. As noted, it is trivial to estimate *p_j_*, and guaranteed that the *p_j_* will be both unique and monotone. To facilitate the discussion of the more general case (where there are zero counts), consider this ideal problem (no zeros) in the context of standard LAD regression. For notational convenience, make the following definitions:

$$SUD_{j}^{'}\left( \beta_{0j} \right):=\sum_{\{i:z_{i}>\left( \beta_{0j}+d_{i} \right)\}} \left[ z_{i}-\left( \beta_{0j}+d_{i} \right) \right]$$

$$SLD_{j}^{'}\left( \beta_{0j} \right):=\sum_{\{i:z_{i}\leq\left( \beta_{0j}+d_{i} \right)\}} \left[ \left( \beta_{0j}+d_{i} \right)-z_{i} \right]$$

where the abbreviations denote “sum of upper deviations” and “sum of lower deviations” respectively across the collection of cells indexed by *i*.

To solve for a percentile given a quantile, the previous regression problem is inverted – and expanded to remove the *p_π_(⋅ )* notation – to find the set

$$\left\{ \pi:\beta_{0j}=\min_{\beta_{0}} \left( \pi SUD_{j}^{'}\left( \beta_{0} \right)+\left( 1-\pi\right)SLD_{j}^{'}\left( \beta_{0} \right) \right) \right\}$$

Note that *SUD_1_^’^=0* and *SLD_n_^’^=0* and that *SUD_j_^’^* (*SLD_j_^’^*) are increasing (decreasing) in *j*. Additionally, a linear interpolation of *SUD_j_^’^* (*SLD_j_^’^*) would have positive (negative) derivatives. Thus, the surface of the convex hull of the set *{(SUD_j_^’^, SLD_j_^’^)}* contains all points within the set. This means that for each *β_0j_*, there exists some unique *π_j_* for which *β_0j_* is the minimizer of the usual, non-inverted, LAD problem. Specifically, a minimizing *π_j_* is one such that the line *π_j_SUD_j_^’^-(1-π_j_)SLD_j_^’^=c* for some constant *c* is a sub-tangent of the linear interpolation of *{(SUD_j_^’^, SLD_j_^’^)}* at the relevant point.

This can be demonstrated as follows: suppose *-π_j_/(1-π_j_)* is a slope in the sub-derivative of the linear interpolation at point *(SUD_j_^’^, SLD_j_^’^)* so that the convex combination of sums of deviations takes some value, *π_j_SUD_j_^’^+(1-π_j_)SLD_j_^’^=b*. Consider then the point *(SUD_j+1_^’^, SLD_j+1_^’^)* and define *(dU,dL)≔(SUD_j+1_^’^,SLD_j+1_^’^)-(SUD_j_^’^,SLD_j_^’^)* where *dU,-dL>0* so the convex combination at *j+1* can be written as *π­_j_(SUD_j_^’^+dU)+(1-π_j_)(SLD_j_^’^+dL)=b+π_j_dU+(1-π_j_)dL*. Since *-π­_j_/(1-π_j_)* is in the sub-derivative, it is the case that

$$-\frac{\pi_{j}}{\left( 1-\pi_{j} \right)}\leq\frac{dL}{dU}\Longrightarrow\pi_{j}dU+\left( 1-\pi_{j} \right)dL\geq0$$

with equality only if *-π_j_/(1-π­_j_)* is the maximal sub-derivative, showing *β_0(j+1)_* is not a solution for the LAD problem given weight *π_j_* excepting only the minimal *π_j_* allowed by the sub-derivative. A similar result holds for point *j-1*.

Therefore, to solve for a percentile *p_j_*, one can consider the derivatives of the sums of deviations parameterized by the intercept *β_0j_*

$$dSUD_{j}^{'}=\frac{d}{d\left( \beta_{0j} \right)}SUD_{j}^{'}=-\left| \{j:z_{j}>\left( \beta_{0j}+d_{j} \right)\} \right| =-\left( j-1 \right)$$

$$dSLD_{j}^{'}=\frac{d}{d\left( \beta_{0j} \right)}SLD_{j}^{'}=\left| \{j:z_{j}\leq\left( \beta_{0j}+d_{j} \right)\} \right|=\left( n-j+1 \right)$$

where the second equalities follow from the uniqueness and ordering of the indices *j*.

Then the slope of one possible subtangent in terms of *π_j_* is

$$-\frac{\pi_{j}}{\left( 1-\pi_{j} \right)}=\frac{dSLD_{j}^{'}}{dSUD_{j}^{'}}=-\frac{\left( n-j+1 \right)}{\left( j-1 \right)}$$

so

$$p_{j}=\pi_{j}=\frac{-\frac{dSLD_{j}^{'}}{dSUD_{j}^{'}}}{1-\frac{dSLD_{j}^{'}}{dSUD_{j}^{'}}}=\frac{\left( n-j+1 \right)}{\left( j-1 \right)+\left( n-j+1 \right)}=\frac{n-j+1}{n}$$

which is the same result as at the beginning of the previous section.

To generalize solving for percentiles *{p_j_}* in the context of cLAD regression, one need only make a few modifications to the above results. First, define

$$SUD_{j}\left( \beta_{0j} \right)=\sum_{z_{i}\geq\left( \beta_{0j}+d_{i} \right)^{*}} \left[ z_{i}-\left( \beta_{0j}+d_{i} \right)^{*} \right]$$

$$SLD_{j}\left( \beta_{0j} \right)=\sum_{z_{i}<\left( \beta_{0j}+d_{i} \right)^{*}} \left[ \left( \beta_{0j}+d_{i} \right)^{*}-z_{i} \right]$$

for the censoring function *(x)^*^*. Then the percentile problem seeks to find sets of a familiar form:

$$\left\{ \pi:\beta_{0j}=\min_{\beta_{0}} \left( \pi SUD_{j}\left( \beta_{0} \right)+\left( 1-\pi\right)SLD_{j}\left( \beta_{0} \right) \right) \right\}$$

with a familiar solution,

$$-\frac{\pi_{j}}{\left( 1-\pi_{j} \right)}=\frac{dSLD_{j}}{dSUD_{j}}$$

assuming the same convexity conditions hold for the set *{(SUD_j_,SLD_j_)}.*

In the presence of censoring, the convexity conditions may not hold. A correction to enforce convexity is discussed later in the section. The main difference between the general result which accommodates censoring and that for LAD regression is in the precise formulation of the derivatives of the sums of deviations.

$$dSUD_{j}=\frac{d}{d\left( \beta_{0j} \right)}SUD_{j}$$

$$=-\left| \left\{ j:z_{j}>\left( \beta_{0j}+d_{j} \right)^{*},d_{j}\geq\beta_{0j}-\log\left( 0.999 \right) \right\} \right|$$

$$dSLD_{j}=\frac{d}{d\left( \beta_{0j} \right)}SLD_{j}$$

$$=\left| \left\{ j:z_{j}\leq\left( \beta_{0j}+d_{j} \right)^{*},d_{j}\geq\beta_{0j}-\log\left( 0.999 \right) \right\} \right|$$

These derivatives do have a similar interpretation to those of the previous section, however. Specifically, up to a sign change, they are the number of observations above/below the regression line under consideration which *also* have sequencing depth above the point where that regression line hits the censoring threshold of *log(0.999)*.

This gives a convenient interpretation to the solution for *p_j_* as well. The solution itself is

$$p_{j}=\frac{dSLD_{j}}{dSLD_{j}-dSUD_{j}}$$

which is simply the empirical percentile from before, but computed on the subset of observations with sequencing depth above the point where the regression line becomes censored. This is consistent with the result from Powell that cLAD regression is equivalent to LAD regression performed on the subset of data for which the probability of censoring is uniformly no greater than the regression percentile *π* and at some covariates the probability of censoring is strictly less than *π*.

It was previously noted that the censored data do not guarantee the convexity conditions on the set of upper and lower deviations as is the case in traditional LAD regression. This can occur stochastically in the lower quantiles when there are few data points from which to estimate the percentiles. This can also occur systematically when the observed expression is correlated with sequencing depth as may occur when sub-populations of cells express in aggregate at different levels.

To correct for both of these issues simultaneously, a monotonicity condition is imposed on the estimated *p_j_*. First, the *p_j_* are computed only on the subset of upper/lower deviations that exist on the edge of the convex hull of *{SUD_j_, SLD_j_}*. Following computation of percentiles on this subset of quantiles, percentiles are adjusted such that differences between adjacent percentiles are bounded above and below. The bounds are as follows:

$$p_{j}-p_{j+1}\geq\frac{\sum_{k} z_{k}>\left( \beta_{0j}+d_{k} \right)^{*}-\sum_{k} z_{k}>\left( \beta_{0j+1}+d_{k} \right)^{*}}{n}$$

$$p_{j}-p_{j+1}\leq p_{j}-\frac{\sum_{k} z_{k}<\left( \beta_{0j+1}+d_{k} \right)^{*}}{n}$$

### S2 Datasets

#### PBMC_Pure

*PBMC68K_Pure* is a partner dataset to PBMC68K (Zheng et al., 2017) produced by fluorescence activated cell sorting (FACS) of peripheral blood mononuclear cells (PBMCs) into 10 cell types and separately sequencing each group. One group was then computationally separated into two resulting in 11 annotated cell-types. These cell-type annotations are considered here as ground truth when evaluating the effects of normalization on downstream clustering, and for increased accuracy, the six most homogenous annotations from visual inspection of within-annotation tSNE plots (van der Maaten and Hinton, 2008; Van Der Maaten, 2014) were subset: CD4+ T Helper2, CD4+/CD25 T Reg, CD4+/CD45RA+/CD25- Naive T, CD4+/CD45RO+ Memory, CD56+ NK, and CD8+/CD45RA+ Naive Cytotoxic.

UMI count matrices and barcode (cell) metadata are available from the GitHub repository associated with the publication: <https://github.com/10XGenomics/single-cell-3prime-paper/tree/master/pbmc68k_analysis>.

#### PBMC5K_Prot

*PBMC5K_Prot* is a dataset of approximately 5 thousand PBMCs sequenced by and available from 10X genomics under the name “5k Peripheral blood mononuclear cells (PBMCs) from a healthy donor with cell surface proteins (v3 chemistry)” and processed under cell ranger version 3.1.0. A panel of 31 surface proteins were sequenced in parallel with the cDNA libraries. We perform unsupervised clustering on the protein abundance estimates to generate pseudo-annotations independently from RNA expression measurements.

#### MaltTumor10K

*MaltTumor10K* is a dataset of approximately 10 thousand cells from a MALT tumor sequenced by and available from 10X genomics under the name “10k Cells from a MALT Tumor - Gene Expression and Cell Surface Protein” and processed under cell ranger version 3.0.0. A panel of 17 surface proteins were sequenced in parallel with the cDNA libraries. We perform unsupervised clustering on the protein abundance estimates to generate pseudo-annotations independently from RNA expression measurements.

#### MouseBrain

*MouseBrain* is a dataset of approximately 9 thousand mouse brain cells sequenced by and available from 10X genomics under the name “9k Brain Cells from an E18 Mouse” and processed under cell ranger version 1.3.0.

#### PBMC68K

*PBMC68K* is a partner dataset to PBMC68K_Pure (Zheng *et al.*, 2017) produced by sequencing approximately 68 thousand PBMCs. In the original paper, pseudo-annotations were generated by computational matching of these cells to the purified lines of PBMC68K_Pure. We, however, treat these as unannotated cells. UMI count matrices are available from 10X genomics under the name “Fresh 68k PBMCs (Donor A)” and processed under cell ranger version 1.1.0.

#### EMT

*EMT* is a dataset of 5,004 sequenced MCF10A mammary epithelial cells induced to undergo spontaneous endothelial to mesenchymal transitions (EMTs) through the cellular detection of neighboring unoccupied space (McFaline-Figueroa *et al.*, 2019). This spatial effect allowed the authors to dissect an inner region a-priori expected to be primarily endothelial cells and an outer region a-priori expected to be primarily mesenchymal cells which were then sequenced separately. The authors produced another dataset of cells activated by TGF-β (denoted TGFB in the barcode metadata), but we consider only the first dataset (denoted Mock in the barcode metadata). Included in the initial publication, the authors describe eight gene sets from the Hallmark collection (Liberzon *et al.*, 2015) which they consider to be significantly enriched for activity during EMT. We take this set of terms as a ground truth for assessing power under a range of normalization techniques: ESTROGEN RESPONSE LATE, ESTROGEN RESPONSE EARLY, P53 PATHWAY, KRAS SIGNALING DN, MYC TARGETS V1, MYC TARGETS V2, PI3K AKT MTOR SIGNALING, and EPITHELIAL MESENCHYMAL TRANSITION. The UMI count matrices and barcode metadata are available on GEO under accession number GSE114687.

#### Dataset processing

For the un-annotated datasets published by 10X (PBMC5K_Prot, MaltTumor10K, MouseBrain, PBMC68K), the UMI count matrices analyzed in this paper were derived from un-filtered gene-barcode matrices. *emptyDrops* (R package: *DropletUtils*; parameters: lower = 20, niters = 160000, test.ambient = TRUE) was used to differentiate empty droplets from barcodes associated with cells. Cellular barcodes were then defined as those with FDR corrected p-values less than 1e-3. For the datasets with surface protein expression (PBMC5K_Prot and MaltTumor10K), cells were additionally filtered to retain only those with a minimum of 100 protein-specific UMIs. This procedure resulted in datasets with 4978 cells (PBMC5K_Prot), 8670 cells (MaltTumor10K), 3756 cells (MouseBrain), and 77249 cells (PBMC68K). Where applicable, surface protein expression was omitted from the rows of UMI count matrices for downstream analysis and testing.

Pseudo-annotations were generated from the datasets with surface protein expression (PBMC5K_Prot and MaltTumor10K) in a manner similar to the unsupervised clustering of all datasets. Surface protein expression was normalized by the method of median ratio (Anders and Huber, 2010). Note: as all surface proteins are generally expected to have cell-type-specific abundances, this normalization is only expected to equalize counts within cell-types meaning that between cell-type calculations of relative abundance are rendered inapplicable; as our purpose is clustering, this is not a problem. Using the Seurat pipeline, protein expression is reduced to 20 dimensions for PBMC5K_Prot (from 29 distinct proteins) and 15 dimensions for MaltTumor10K (from 17 distinct proteins). Graph based clustering is then performed using *FindNeighbors* and *FindClusters* (additional parameters: algorithm = 3, n.start = 100, n.iter = 100) from the Seurat package.

### S3 Data simulation

#### Initial grouping

We generate our simulated datasets from experimentally derived UMIs with the purpose of making our simulations as representative of the characteristics of observed data as possible. The first step is to normalize experimental data using Dino (see S4.2) and then perform unsupervised clustering on the normalized data (see S4.4). This results in (relatively) homogenous subset of cells. We then generate individual clusters of simulated data from each of these discovered clusters of experimental data, which are merged into a heterogenous test dataset.

#### Group filtering

We vary the size of simulated clusters by powers of 2 (40 cells, 80 cells, 160 cells, etc.). To this end, we calculate the largest *k* such that we have *k* discovered clusters with at least *40×2^k^* cells and discard the remaining experimental data. If *k=6*, then we have exactly 6 clusters of experimental data with at least *40×2^6^=2560* cells, and we discard cells from any smaller clusters.

#### Cluster pair simulation

The simulated dataset is constructed of pairs of simulated clusters for which the EE and DE genes are known. In the case were *k=6* as above, the simulated datasets then consist of 6 cluster pairs, or 12 simulated clusters total. Within a cluster pair, there is an induced difference in sequencing depth and DE genes are randomly selected. Between cluster pairs, there may also be systematic differences in sequencing depth, but only to the extent that there are systematic differences in the depths of the experimental cells these cluster pairs are based on.

To construct one cluster pair, an experimental cluster, denoted by *C_k_*, is randomly sampled. Each of the two simulated clusters in the pair will consist of 40 cells if this is the first cluster pair, 80 cells if this is the second cluster pair, and so on increasing by factors of 2. Denote the number of simulated cells in each cluster by *n*. To generate the first *n/2* cells in each cluster, we sample *n* cells from the experimental data. In order of increasing depth, we sum pairs of experimental cells to create *n/2* pseudo-cells with roughly double the sequencing depth of either of the cells they are comprised of.

Denote the simulated clusters in the pair by *A* and *B*, each to consist of *n* simulated cells. To induce a difference in sequencing depth between *A* and *B*, we sample a sequencing depth fold change, *δ_fc_*, from the range *3/*2 to *4*, and for convenience assign *A* to be the higher depth group. The first *n/2* cells in each group will be generated by binomial sampling from the *n/2* pseudo-cells, with the induced fold change in depth arising from differences in the binomial probability parameter, *p*. Some algebra shows that the choice of

$$p=0.5\pm\frac{(\delta_{fc}-1)}{2(\delta_{fc}+1)}$$

for *A* and *B* respectively will produce clusters with all EE genes once sequencing depth is accounted for under normalization. Specifically, for pseudo-cell *s_j_* and simulated cells *a_j_* and *b_j_* from *A* and *B*, we simulated as

$$a_{j}\sim Binom\left( s_{j},p_{+} \right)$$

$$b_{j}\sim Binom(s_{j},p_{-})$$

where *p_+_* and *p_-_* are the two variants of *p* respectively.

However, this approach only generates EE genes. To simulate known DE genes, we subset those genes in *C_k_* with at least 25% non-zeros. From this set of genes, we sample 10 to be induced DE genes, with sampling weighted by the inverse density of log gene expression, calculated simply as the log of the mean UMIs in *C_k_*. As with the fold change in depth, we sample 10 DE fold changes from the range *3/2* to *6*, denoted by *γ_fc,g_* with the subscript *g* indexing the 10 gene-specific DE fold changes. As we do not want all DE genes to be upregulated in *A*, we invert each of the *γ_fc,g_* with probability 0.5. If we now consider the binomial probability, *p*, to be a vector of length equal to the number of genes, and *p_DE_* to denote the subset of elements which are DE after correcting for sequencing depth, some similar algebra to the above shows that

$$p_{DE, g}=0.5\pm\frac{(\delta_{fc}\gamma_{fc,g}-1)}{2(\delta_{fc}\gamma_{fc,g}+1)}$$

where this formulation also allows the definition of *a_j_* and *b_j_* to be defined as the same binomial random variable parameterized by *p_+_* or *p_-_*, where *p* now includes information about DE sampling.

Two problems remain to be addressed; that we have only discussed the generation of *n/2* of the cells in each group and that correcting for sequencing depth as defined here can induce slight but systematic differential expression in the EE genes. Take, for example, the extreme case where all the DE genes are upregulated in *A* relative to *B*, after correcting for sequencing depth. In this case, calculating sequencing depth from the sum of simulated UMIs within a cell, and correcting for that depth, will induce a slight but consistent down-regulation in the EE genes in *A* relative to *B*. We address this by adding a correction factor to the remaining *n/2* simulated cells in each cluster.

The degree of this induced bias can be simply calculated as the ratio of expected total sequencing depth (total meaning summed across cells as well as genes) under the above DE model and the model where all genes are simulated EE. Let *p_+_* be, as above, a vector of binomial probabilities which includes DE information for the simulation of *A* and let *p_EE+_* be a corresponding binomial probability vector for which all genes are EE, that is, suppose all elements of *p_EE+_* are equal to the original, scalar, definition of *p_+_*. Let *C_k-_* denote a vector of gene-wise UMIs, summed across all cells in *C_k_*. Then, the total expected depth for the EE case is *p_EE+_^T^C_k-_* and the degree of bias in the above simulated cells, inducing DE in simulated EE genes, is

$$\alpha_{+bias}=\frac{p_{+}^{T}C_{k-}}{p_{EE+}^{T}C_{k-}}$$

This can be interpreted as implying that normalized EE genes in *A* will, on average, be a factor of *1/α_+bias_* different from what would have been the case had all genes had been simulated as EE. If *α_-bias_=α_+bias_*, then this wouldn’t be a problem, but such is not the case. Unfortunately, it is also the case that *α_-bias_≠1/α_+bias_*, as can be shown by simple counter examples. Therefore, we compute separate corrective factors for *A* and *B* under the principle that expression of EE genes between *A* and *B* should, when averaged across the first *n/2* cells and the second, corrected *n/2* cells demonstrate the desired fold change in depth. This leads to the corrective factor, *c*

$$\left( \frac{c_{+}p_{+}^{T}C_{k-}}{p_{EE}^{T}C_{k-}} \right)^{-1}=1-\left( \frac{1}{\alpha_{+bias}}-1 \right)$$

$$\Rightarrow c_{+}=\frac{1}{2\alpha_{+bias}-1}$$

$$\Rightarrow c_{-}=\frac{1}{2\alpha_{-bias}-1}$$

This then fully defines the simulated cells:

$$a_{j}\sim\left\{ \begin{aligned} Binom\left( s_{j},p_{+} \right), j\leq n/2 \\ Binom\left( s_{j-\frac{n}{2}}, c_{+}p_{+} \right), j>n/2 \end{aligned} \right.$$

$$b_{j}\sim\left\{ \begin{aligned} Binom\left( s_{j},p_{-} \right), j\leq n/2 \\ Binom\left( s_{j-\frac{n}{2}}, c_{-}p_{-} \right), j>n/2 \end{aligned} \right.$$

To complete a simulated dataset, the above steps for generating the cluster pair *A* and *B* are repeated for the remaining experimental clusters, generating a heterogenous samples of simulated data for which pairs of simulated clusters have known EE and DE genes.

### S4 Implementation details

#### R package versions

BiocParallel (v1.22.0), BiocSingular (v1.4.0), Dino (v0.6.1), DropletUtils (v1.8.0), irlba (v2.3.3), MAST (v1.14.0), Matrix (v1.2-18), matrixStats (0.56.0), mclust (v5.4.6), monocle (v2.16.0), piano (v2.4.0), snowfall (v1.84-6.1), Scran (v1.16.0), sctransform (v0.2.1), Seurat (v3.2.0)

#### Normalization defaults

**UMI:** reference gene-by-barcode matrix of unique molecular identifiers (UMIs)

**TPM:** rescaling of UMI such that each column (cell or barcode) sums to one million

**TPT:** rescaling of UMI such that each column sums to ten thousand

**Scran:** rescaling of UMI such that each column is divided by the scale factor computed by the scran method from the sizeFactors function in the scran package (Lun *et al.* 2016). Default parameters are used.

**scTrans:** normalized gene-by-barcode matrix output by the vst function in the sctransform package (Hafemeister and Satija, 2019). Default parameters are used excepting: return_cell_attr = TRUE, res_clip_range = c(-50, 50).

**scTransCnt:** corrected UMI count matrix output by the correct_counts function from the sctransform package. Default parameters are used with scTrans and UMI as input.

**Dino:** normalized gene-by-barcode matrix output by the Dino function in the Dino package. Default parameters are used excepting: nCores = 4.

#### Pseudo-time differential expression

Analysis of the EMT dataset in a manner similar to that of the original authors is primarily conducted using Monocle2 (in the monacle package) for pseudo-time ordering and DE testing; piano is used for term enrichment testing. Normalized matrices are used to create monocle objects, and are log transformed with a +1 pseudo count for variance stabilization excepting scTrans which performs variance stabilization internally to the algorithm. For each normalization method, the top 1,000 highest variance genes (after the above-mentioned transformation) are used to perform dimension reduction in the Monocle2 environment which includes construction of a minimum spanning tree using the DDRTree algorithm. This tree is used to construct pseudo-times for each cell, again using the default Monocle2 pipeline. To root the tree (determine which cell is assigned the pseudo-time of 0), we subset the cells assigned to the branches that contain the default earliest and latest pseudo-time, which denote ends of the longest contiguous differentiation path in well-ordered data. Of these two branches, we denote the branch with the highest proportion of inner section cells (expected to be primarily endothelial cells) as the root of the tree. Unlike the analysis in the original paper, trees constructed in our application included branches. Because of this, we remove from further analysis cells in any branches not along the main path, defined as the minimum path between the earliest and latest branches (mentioned in the content on rooting). As such, this sub-tree will be a non-branching path.

Following computation of pseudo-times and sub-setting of cells, we conduct DE tests, here defined as a change in expression over time. Formally, this test is a likelihood ratio test between a natural spline with three degrees of freedom regressed against pseudo-time and an intercept-only model. This test (function differentialGeneTest) is part of the Monocle2 pipeline and was implemented using default parameters.

In their original paper, the authors defined a list of hallmark gene sets as enriched for expression changes over the endothelial to mesenchymal transition; we consider these as ground truth. Enrichment testing for the hallmark collection of gene sets is performed using the piano package (runGSA function) using DE test p-values as gene-level statistics and the “tailStrength” statistical test.

#### Unsupervised clustering

Unsupervised clustering is performed using standard functions in the Seurat package. Normalized matrices are used to create Seurat objects, and are log transformed with a +1 pseudo count for variance stabilization excepting scTrans which performs variance stabilization internally to the algorithm. The top 1,000 highest variance genes are subset to perform dimension reduction using approximate PCA down to 20 dimensions. Per-cell cluster memberships are identified using graph-based clustering using the FindClusters function in Seurat with default parameters excepting: algorithm = 3, n.start = 50, n.iter = 50.

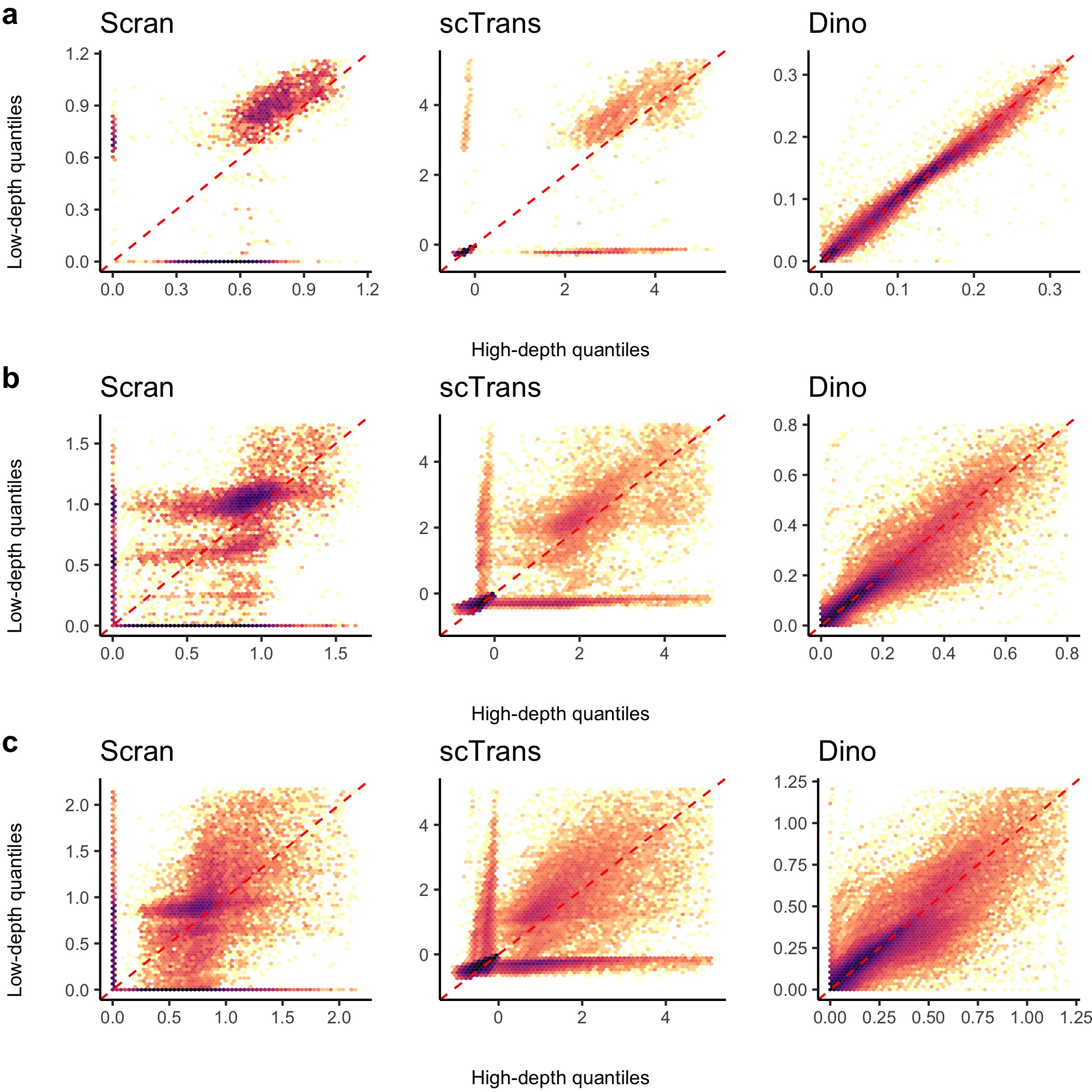

Supplementary Figure S1: **Evaluation of normalized expression distributions across genes.** Quantile-quantile density plots comparing expression quantiles in the high-depth (x-coordinate) and low-depth (y-coordinate) cells across genes and cell-type annotations are shown for the PBMC68K_Pure dataset (a), the MaltTumor10K dataset (b), and the PBMC5K_Prot dataset (c). Colors are log-scale and datapoints in a small neighborhood of zero are omitted to improve visualization.

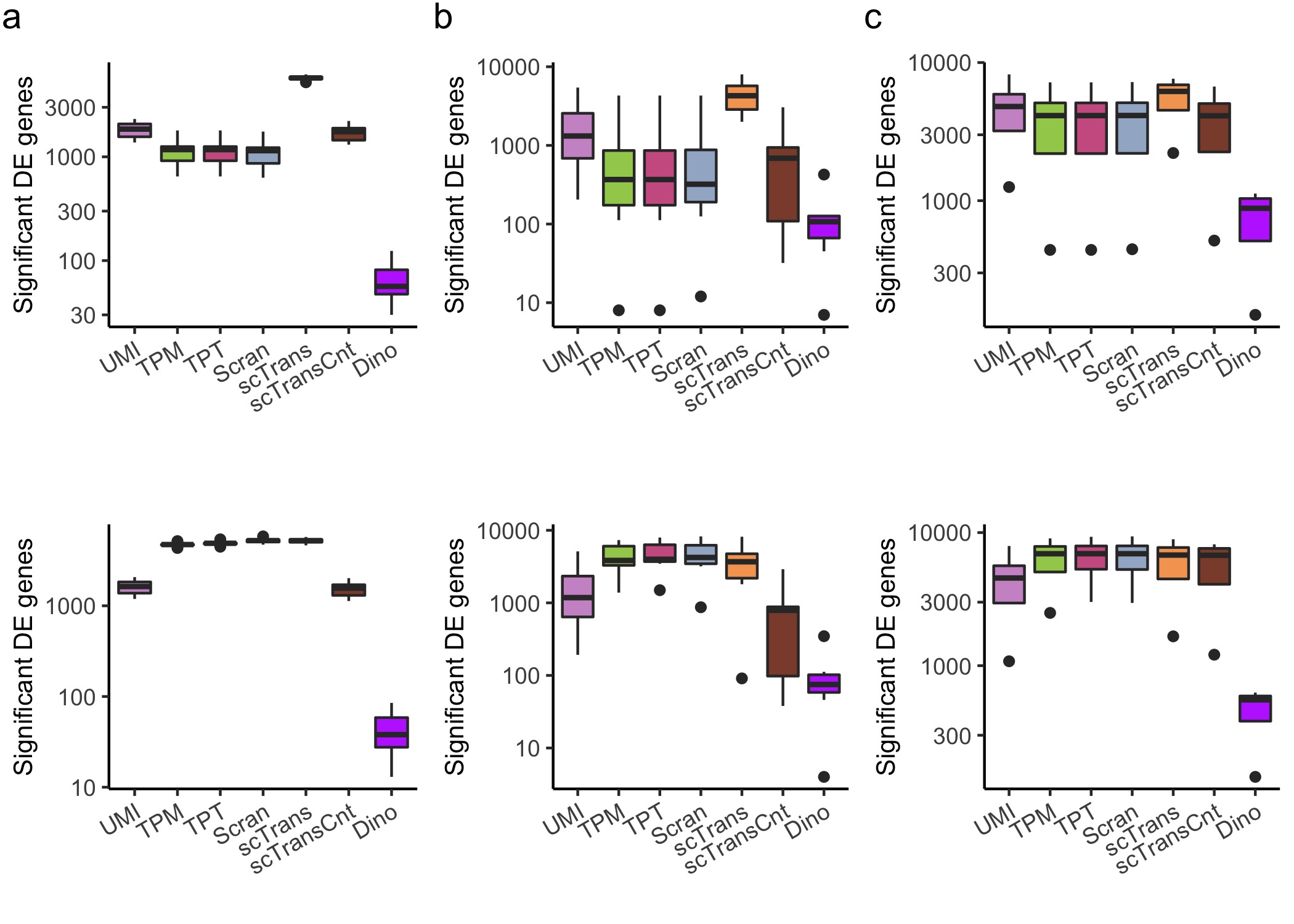

Supplementary Figure S2: **The effects of normalization on downstream DE testing false positives.** Expression data were normalized and genes were tested for DE using a Wilcoxon rank sum test (top) and a MAST test (bottom) between low-depth and high-depth cells within cell-type annotations. Box plots show DE genes in the PBMC68K_Pure dataset (a), the MaltTumor10K dataset (b), and the PBMC5K_Prot dataset (c). Given that cells only differ in depth, DE identifications are expected to be false positives.

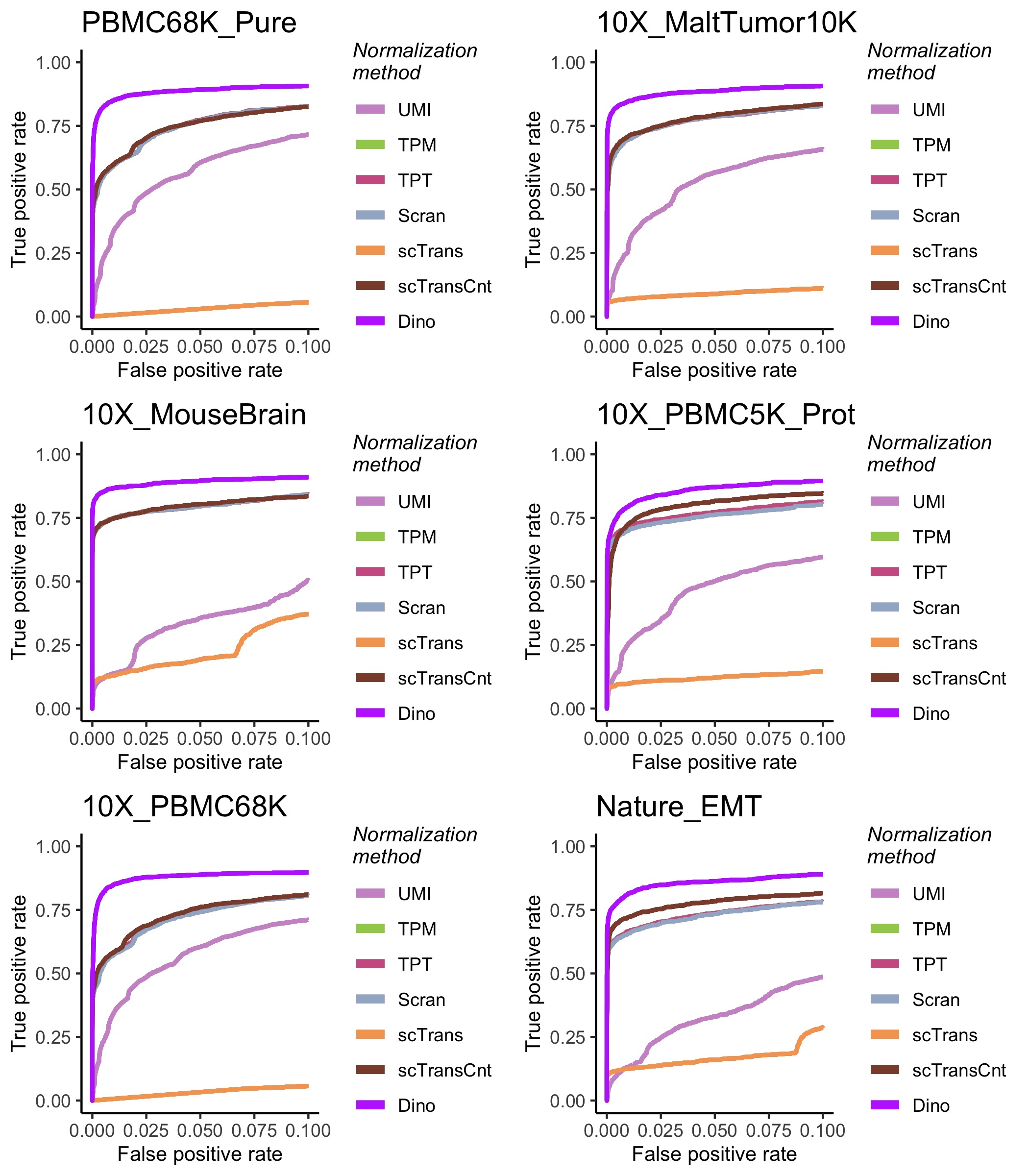

Supplementary Figure S3: **The effects of normalization on downstream DE analysis.** Simulated data based on each of the considered datasets were normalized using each method. ROC curves colored by normalization method define the relationship between average TPR (Power) and average FPR for a Wilcoxon rank sum test, where the average is calculated across 50 simulations from each dataset.

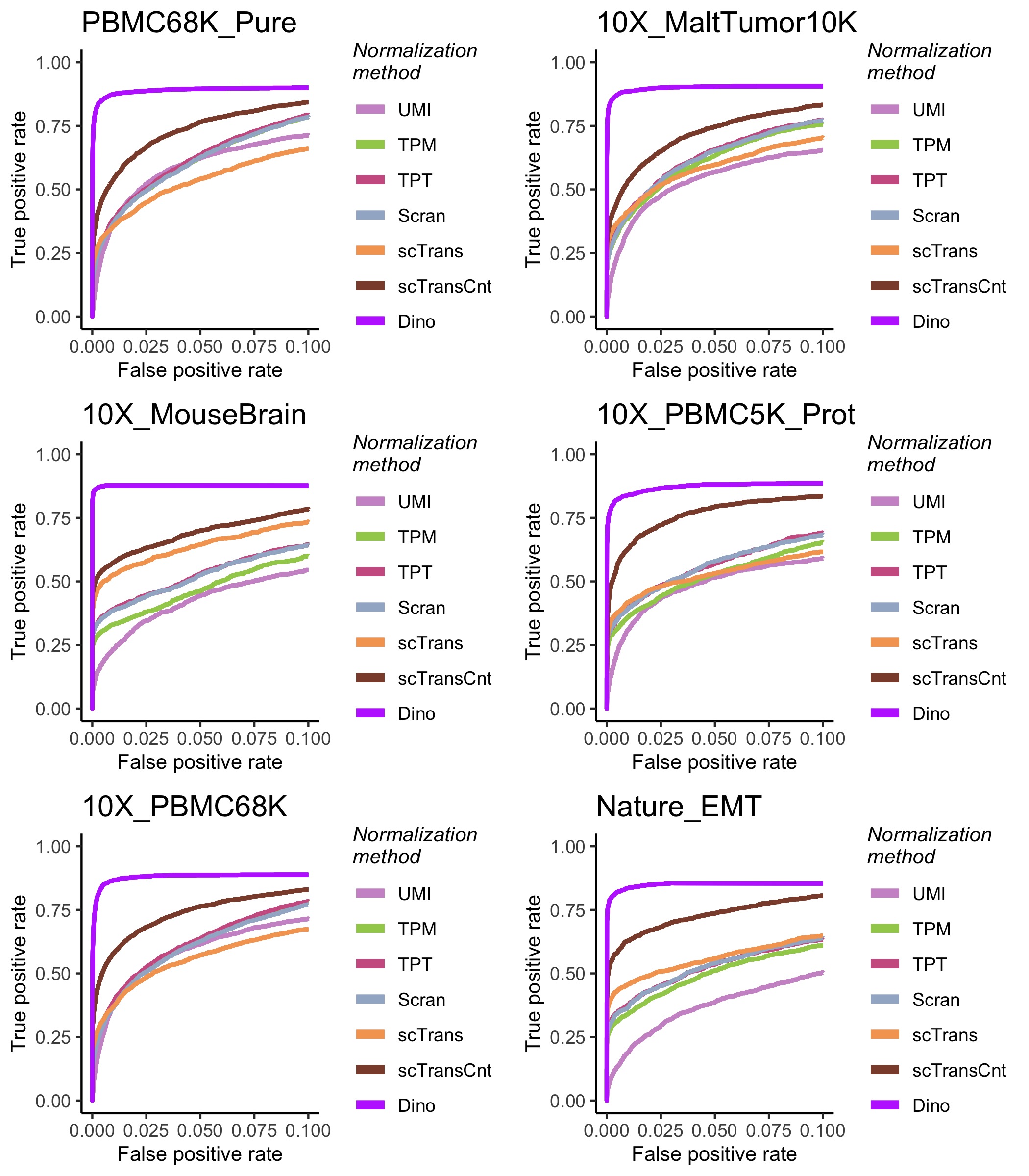

Supplementary Figure S4: **The effects of normalization on downstream DE analysis.** Simulated data based on each of the considered datasets were normalized using each method. ROC curves colored by normalization method define the relationship between average TPR (Power) and average FPR for a MAST test, where the average is calculated across 50 simulations from each dataset.

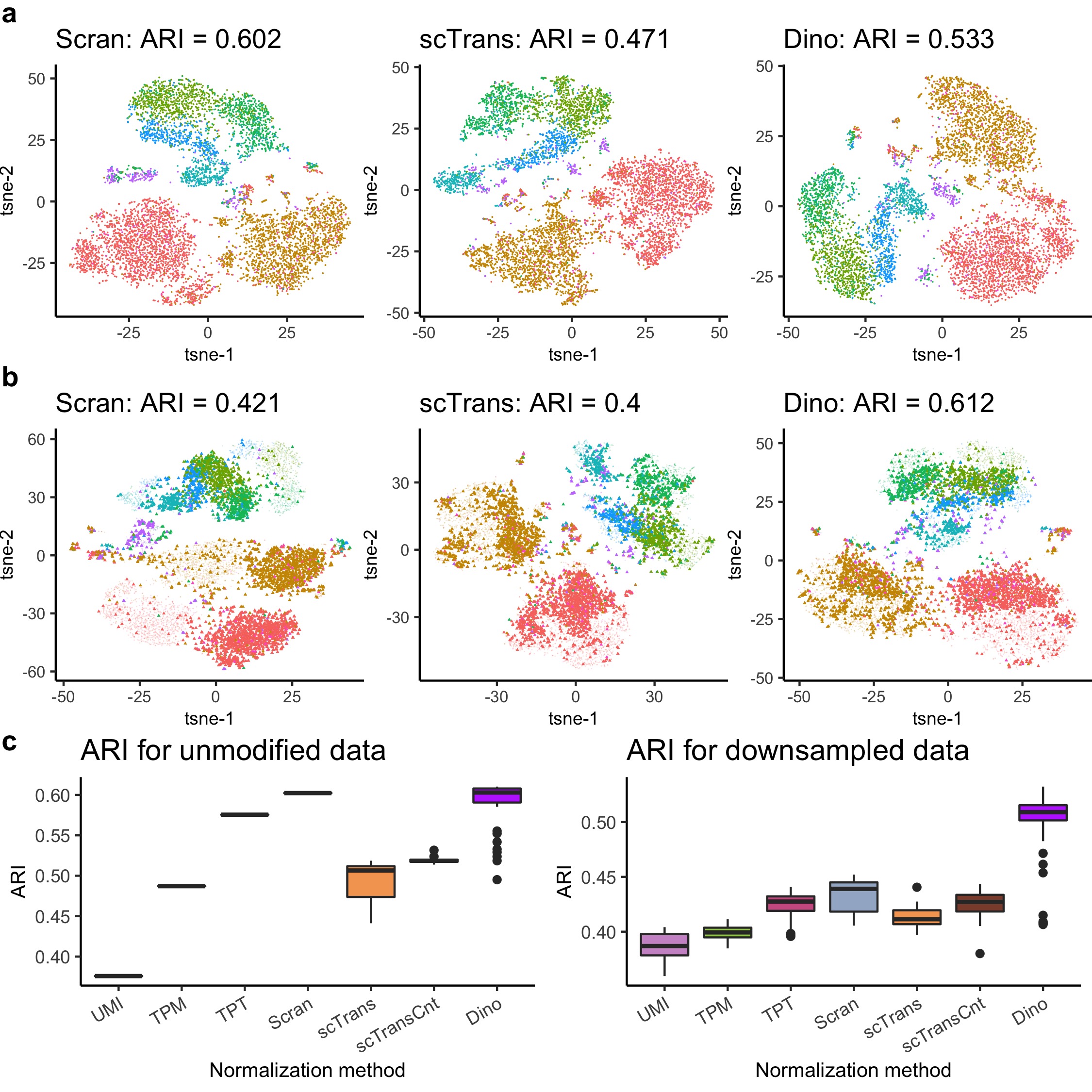

Supplementary Figure S5: **The effects of normalization on clustering.** a) tSNE plots of normalized MaltTumor10K data, colored by 11 cell-type annotations, show similarly high accuracy between methods. b) The same clustering plots as in (a), but with half the data down-sampled prior to normalization to produce greater differences in sequencing depths. c) Boxplots of ARIs for multiple un-modified and down-sampled datasets.

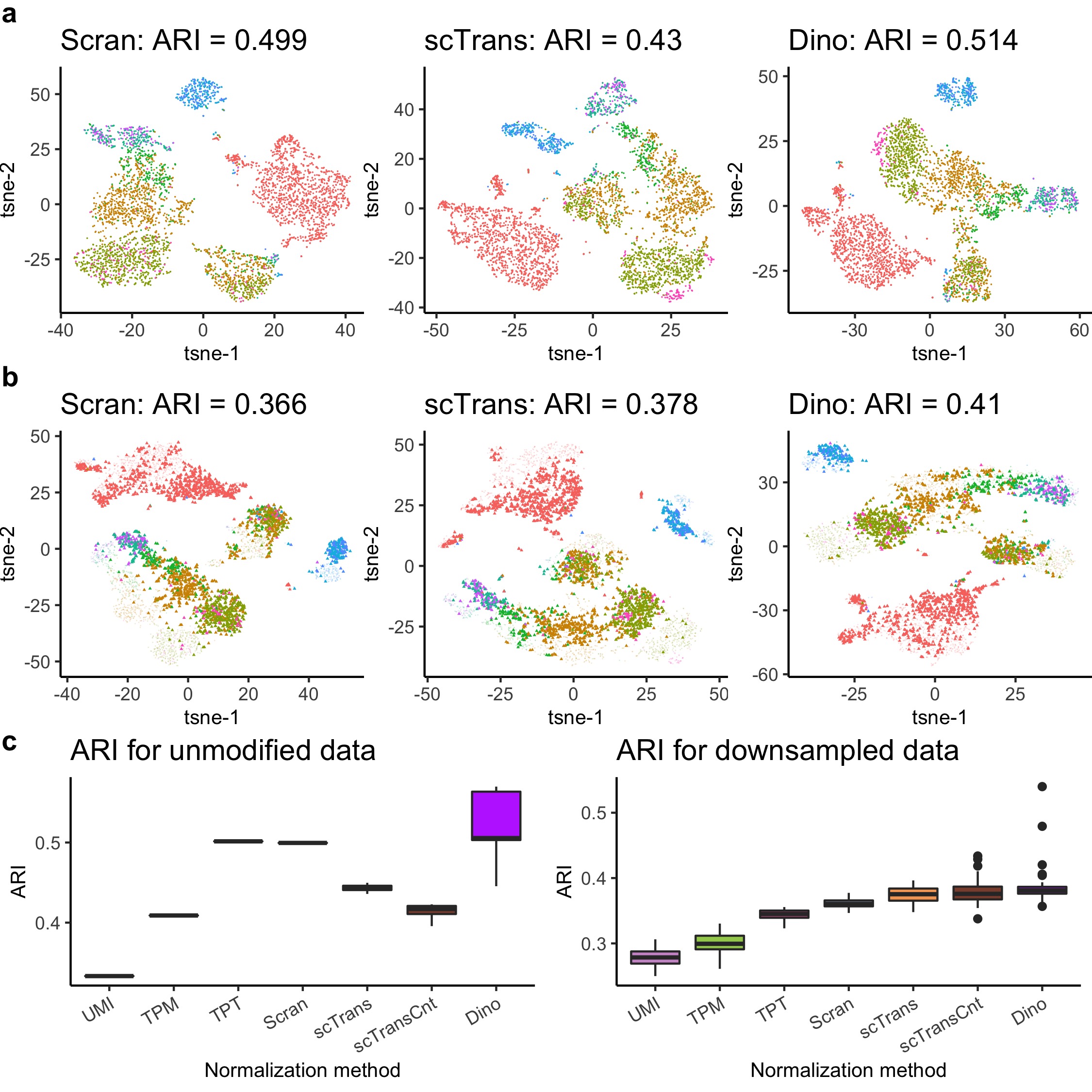

Supplementary Figure S6: **The effects of normalization on clustering.** a) tSNE plots of normalized PBMC5K_Prot data, colored by 13 cell-type annotations, show similarly high accuracy between methods. b) The same clustering plots as in (a), but with half the data down-sampled prior to normalization to produce greater differences in sequencing depths. c) Boxplots of ARIs for multiple un-modified and down-sampled datasets.

Supplementary Table S1: **Average power and FPR across simulated datasets and normalization methods.** 50 simulated datasets are produced from each case study dataset. In each simulation, the data are normalized by the panel of methods and DE genes are identified using the Wilcoxon test. DE genes are defined as those with a Benjamini and Hochberg adjusted p-value less than 0.01. Average power and FPR is calculated over the 50 simulated datasets (standard error computed across simulations).

| **Norm. Method** | ***UMI*** | ***TPT*** | ***TPM*** | ***Scran*** | ***scTrans*** | ***scTransCnt*** | ***Dino*** |  |
| --- | --- | --- | --- | --- | --- | --- | --- | --- |
| **PBMC68K_Pure** | 0.792 (0.007) | 0.862 (0.006) | 0.862 (0.006) | 0.861 (0.006) | 0.902 (0.006) | 0.848 (0.005) | 0.832 (0.006) | **Power** |
|  | 0.199 (0.005) | 0.155 (0.006) | 0.155 (0.006) | 0.155 (0.006) | 0.625 (0.021) | 0.147 (0.005) | 0.006 (0.001) | **FPR** |
| **PBMC5K_Prot** | 0.725 (0.011) | 0.802 (0.009) | 0.802 (0.009) | 0.79 (0.01) | 0.835 (0.009) | 0.785 (0.008) | 0.754 (0.009) | **Power** |
|  | 0.248 (0.013) | 0.087 (0.012) | 0.087 (0.012) | 0.087 (0.012) | 0.62 (0.017) | 0.026 (0.005) | 0.005 (0.001) | **FPR** |
| **MaltTumor10K** | 0.742 (0.009) | 0.833 (0.008) | 0.833 (0.008) | 0.832 (0.007) | 0.896 (0.007) | 0.832 (0.007) | 0.799 (0.009) | **Power** |
|  | 0.213 (0.008) | 0.107 (0.01) | 0.107 (0.01) | 0.106 (0.01) | 0.704 (0.02) | 0.095 (0.008) | 0.002 (0) | **FPR** |
| **MouseBrain** | 0.806 (0.012) | 0.85 (0.01) | 0.85 (0.01) | 0.851 (0.01) | 0.9 (0.007) | 0.854 (0.01) | 0.784 (0.011) | **Power** |
|  | 0.477 (0.016) | 0.112 (0.014) | 0.112 (0.014) | 0.112 (0.014) | 0.507 (0.006) | 0.131 (0.013) | 0 (0) | **FPR** |
| **PBMC68K** | 0.776 (0.008) | 0.842 (0.008) | 0.842 (0.008) | 0.836 (0.008) | 0.879 (0.006) | 0.826 (0.006) | 0.814 (0.007) | **Power** |
|  | 0.181 (0.006) | 0.147 (0.007) | 0.147 (0.007) | 0.146 (0.007) | 0.591 (0.017) | 0.129 (0.008) | 0.005 (0) | **FPR** |
| **EMT** | 0.732 (0.011) | 0.777 (0.011) | 0.777 (0.011) | 0.774 (0.011) | 0.875 (0.009) | 0.789 (0.009) | 0.729 (0.015) | **Power** |
|  | 0.366 (0.013) | 0.09 (0.012) | 0.09 (0.012) | 0.092 (0.012) | 0.56 (0.007) | 0.054 (0.007) | 0.001 (0) | **FPR** |

Supplementary Table S2: **Average power and FPR across simulated datasets and normalization methods.** 50 simulated datasets are produced from each case study dataset. In each simulation, the data are normalized by the panel of methods and DE genes are identified using the MAST test. DE genes are defined as those with a Benjamini and Hochberg adjusted p-value less than 0.01. Average power and FPR is calculated over the 50 simulated datasets (standard error computed across simulations).

| **Norm. Method** | ***UMI*** | ***TPT*** | ***TPM*** | ***Scran*** | ***scTrans*** | ***scTransCnt*** | ***Dino*** |  |
| --- | --- | --- | --- | --- | --- | --- | --- | --- |
| **PBMC68K_Pure** | 0.781 (0.008) | 0.946 (0.005) | 0.944 (0.005) | 0.959 (0.005) | 0.91 (0.006) | 0.895 (0.005) | 0.836 (0.007) | **Power** |
|  | 0.175 (0.008) | 0.272 (0.005) | 0.267 (0.005) | 0.299 (0.005) | 0.428 (0.006) | 0.175 (0.005) | 0.003 (0.007) | **FPR** |
| **PBMC5K_Prot** | 0.701 (0.012) | 0.904 (0.011) | 0.912 (0.011) | 0.932 (0.008) | 0.813 (0.009) | 0.805 (0.008) | 0.753 (0.008) | **Power** |
|  | 0.217 (0.012) | 0.346 (0.011) | 0.348 (0.011) | 0.371 (0.008) | 0.365 (0.009) | 0.06 (0.008) | 0.001 (0.008) | **FPR** |
| **MaltTumor10K** | 0.722 (0.009) | 0.945 (0.006) | 0.943 (0.006) | 0.956 (0.005) | 0.893 (0.008) | 0.893 (0.007) | 0.804 (0.009) | **Power** |
|  | 0.185 (0.009) | 0.324 (0.006) | 0.31 (0.006) | 0.338 (0.005) | 0.38 (0.008) | 0.164 (0.007) | 0.001 (0.009) | **FPR** |
| **MouseBrain** | 0.781 (0.011) | 0.957 (0.005) | 0.975 (0.004) | 0.973 (0.004) | 0.891 (0.008) | 0.902 (0.009) | 0.786 (0.011) | **Power** |
|  | 0.437 (0.011) | 0.54 (0.005) | 0.584 (0.004) | 0.572 (0.004) | 0.3 (0.008) | 0.262 (0.009) | 0 (0.011) | **FPR** |
| **PBMC68K** | 0.766 (0.008) | 0.933 (0.006) | 0.931 (0.006) | 0.947 (0.006) | 0.894 (0.006) | 0.858 (0.007) | 0.814 (0.007) | **Power** |
|  | 0.16 (0.008) | 0.248 (0.006) | 0.247 (0.006) | 0.282 (0.006) | 0.369 (0.006) | 0.137 (0.007) | 0.003 (0.007) | **FPR** |
| **EMT** | 0.71 (0.011) | 0.925 (0.01) | 0.94 (0.008) | 0.934 (0.009) | 0.847 (0.01) | 0.819 (0.01) | 0.721 (0.013) | **Power** |
|  | 0.332 (0.011) | 0.474 (0.01) | 0.481 (0.008) | 0.465 (0.009) | 0.321 (0.01) | 0.124 (0.01) | 0 (0.013) | **FPR** |
